# ProSAAS is Preferentially Secreted from Neurons During Homeostatic Scaling and Reduces Amyloid Plaque Size in the 5xFAD Mouse Hippocampus

**DOI:** 10.1101/2024.04.18.590133

**Authors:** Samira Mitias, Nicholas Schaffer, Saaya Nair, Chelsea Hook, Iris Lindberg

## Abstract

The accumulation of β-amyloid in Alzheimer’s disease greatly impacts neuronal health and synaptic function. To maintain network stability in the face of altered synaptic activity, neurons engage a feedback mechanism termed homeostatic scaling; however, this process is thought to be disrupted during disease progression. Previous proteomics studies have shown that one of the most highly regulated proteins in cell culture models of homeostatic scaling is the small secretory chaperone proSAAS. Our prior work has shown that proSAAS exhibits anti-aggregant behavior against alpha synuclein and β-amyloid fibrillation *in vitro*, and is upregulated in cell models of proteostatic stress. However, the specific role that this protein might play in homeostatic scaling, and its anti-aggregant role in Alzheimer’s progression, is not clear. To learn more about the role of proSAAS in maintaining hippocampal proteostasis, we compared its expression in a primary neuron model of homeostatic scaling to other synaptic components using Western blotting and qPCR, revealing that proSAAS protein responses to homeostatic up- and down-regulation were significantly higher than those of two other synaptic vesicle components, 7B2 and carboxypeptidase E. However, proSAAS mRNA expression was static, suggesting translational control (and/or reduced degradation). ProSAAS was readily released upon depolarization of differentiated hippocampal cultures, supporting its synaptic localization. Immunohistochemical analysis demonstrated abundant proSAAS within the mossy fiber layer of the hippocampus in both wild-type and 5xFAD mice; in the latter, proSAAS was also concentrated around amyloid plaques. Interestingly, overexpression of proSAAS in the CA1 region via stereotaxic injection of proSAAS-encoding AAV2/1 significantly decreased amyloid plaque burden in 5xFAD mice. We hypothesize that dynamic changes in proSAAS expression play a critical role in hippocampal proteostatic processes, both in the context of normal homeostatic plasticity and in the control of protein aggregation during Alzheimer’s disease progression.

## Introduction

Alzheimer’s disease (AD) is the most prominent form of dementia in the elderly and is a growing health concern due to increased aging of the US population, increasing health care costs; and the lack of therapeutic options. While the cellular and molecular mechanisms underlying disease progression are under active investigation, the accumulation of β-amyloid (Aβ)-containing plaques within the brain is a recognized hallmark of AD, and there is increasing evidence that soluble Aβ oligomers induce synaptic damage and impairment of homeostatic plasticity (1,2).

Homeostatic plasticity is a widely accepted mechanism for long-term activity-dependent alterations in synaptic strength, and encompasses both long-term potentiation (LTP), and its analog, long-term depression (LTD) (3). Strong activity in both pre- and post-synaptic neurons is essential for reinforcement of these synapses; however, induced activity changes pose potential problems in terms of runaway excitation or uncontrolled synaptic pruning (reviewed in (4)). To prevent hyperactivity and uncontrolled synaptic refinement, neurons engage a negative feedback mechanism termed homeostatic synaptic plasticity to maintain neuronal activity within a physiological range. Homeostatic plasticity is thought to rely on the synthesis and degradation of a variety of proteins involved in vesicle trafficking, neurotransmitter release, and clearance of the synaptic cleft (5).

Homeostatic mechanisms become defective in neurons expressing AD-related transgenes, as these neurons are unable to compensate for disruptions in network activity (6). The inability of neurons to constrain neuronal firing within a physiological limit leads to excessive strengthening or weakening of synapses, and eventually, neurological disorders (reviewed in (4)). Thus, the identification of the molecular players involved in synaptic plasticity is important not only for gaining insight into the maintenance of normal network activity, but for understanding homeostatic disturbances resulting from disease progression.

Dynamic changes in synaptic proteins during homeostatic scaling are likely to contribute greatly to synaptic homeostasis. In a proteomics study by Dörrbaum *et al.* (7), the neuronal secretory chaperone proSAAS (gene name *Pcsk1n*) was found to be the most highly upregulated protein in a drug-induced homeostatic scaling model. The small proSAAS protein, named for a four-residue segment present in the mammalian protein, is widely distributed within the brain and endocrine systems (8) and is associated with maintaining proteostasis upon endoplasmic reticulum stress (9). ProSAAS overexpression also protects cells from cytotoxic oligomers associated with various neurodegenerative diseases, including those found in Parkinson’s disease (10,11) as well as in Alzheimer’s disease (12,13). The potent anti-aggregation properties of proSAAS are likely attributable to its intrinsically disordered secondary structure, which confers unique biochemical properties (14). However, it is not yet clear why this protein, whose synaptic functions are as yet undefined, consistently appears at the top of the proteomics lists during homeostatic scaling experiments (7,15).

To validate proteomics findings and obtain further information on proSAAS changes during homeostatic scaling, we have performed RNA and protein measurements in differentiated hippocampal cultures subjected to homeostatic scaling and have examined proSAAS distribution in normal and diseased brain sections. Our work confirms that proSAAS indeed exhibits disproportionate homeostatic changes compared to other synaptic components. We also show that proSAAS is highly expressed in the mossy fibers of the dentate gyrus and surrounds amyloid plaques in an AD model mouse, suggesting that it serves a potentially important function in maintaining synaptic health.

## Materials and Methods

### Materials

Bicuculline (BIC), tetrodotoxin (TTX), and muscimol were obtained from Sigma (St. Louis, MO). Oligonucleotides were synthesized by Integrated DNA Technologies (Coralville, IA). AAV2/1 viruses, with proSAAS or GFP driven by the human synapsin promoter, were produced and purified by the University of Maryland Virus Core Facility.

### Primary Neuron Cultures

Surplus primary neurons, prepared by the Blanpied laboratory (University of Maryland, Baltimore) from the pooled hippocampi of rat embryos, were used in this work. Briefly, the hippocampi from approximately 20 E17-E18 embryos were minced into small pieces in sterile dissection media (Hanks’ balanced salt solution (Gibco; Waltman, MA), 10 mM HEPES, 33.3 mM glucose, 5 μg/ml gentamycin, 0.3% bovine serum albumin, and 2 mM MgSO4, pH 7.4), digested in 0.25% trypsin-EDTA, and mechanically dissociated by pipetting through a series of Pasteur pipettes with different bore sizes, as previously described (16).

For drug treatment experiments, 250,000 hippocampal cells were plated into a 12-well poly-l-lysine-coated plate in Neurobasal Medium (Invitrogen; Waltman, MA) supplemented with 1 μg/ml gentamycin (VWR; Radnor, PA; #97061-370), 2 mM GlutaMAX (Invitrogen #35050061), 5% B27 supplement (Gibco) and 5% Fetal Bovine Serum (FBS) (Gemini Bio; West Sacramento, CA; #100-106-500). Serum was removed in a medium change the next day. Dividing cells were eliminated by the inclusion of fluorodeoxyuridine at 0.2 μM on Day 3 *in vitro*. Neurons were differentiated for the following 10-14 days, with twice weekly partial medium changes, in a humidified incubator maintained at 37°C and 5% CO2. For secretion experiments, 1 x 10^6^ hippocampal cells were plated into each well of a 6-well poly-l-lysine coated plate to ensure detectable protein levels in the medium.

### Drug Treatment of Primary Hippocampal Neurons

Hippocampal cultures were treated with TTX (2 μM final concentration), BIC (20 μM), or the acetic acid vehicle (1 mM) in fresh media. After 24 h of incubation in the presence of the respective drug or vehicle control, hippocampal cells were washed with Dulbecco’s phosphate-buffered saline (dPBS; Invitrogen), then solubilized in 250 μl Laemmli sample buffer containing 6 M urea (50 mM Tris-HCl pH 6.8, 10% glycerol, 2% SDS, 5% beta mercaptoethanol, 6 M urea, 0.8% Bromophenol Blue). Samples were immediately heated at 95°C for 5 min, then sonicated for 20 sec to reduce viscosity. For time course experiments, hippocampal cells were harvested at 12, 24, 48, and 72 h after drug addition.

For secretion experiments, following 24 h of 2 μM TTX treatment of hippocampal cells, media were removed and replaced with warmed OptiMEM (Invitrogen) containing 0.1% aprotinin for 30 min to equilibrate cells in the new medium. This medium was then replaced with OptiMEM containing 0.1% aprotinin for 2 h, and this was collected as “basal secretion medium”. Cells were then treated with 45 mM KCl in OptiMEM with 0.1% aprotinin for 2 h and this was collected as “stimulated secretion medium”. Two volumes of ice-cold methanol were added to all medium samples along with 2 μg of carrier glycogen, and samples left on ice for 1 h. Mixtures were centrifuged at 14,000 rpm in the cold, supernatants were removed, and pellets were resuspended in 50 μl of Laemmli sample buffer containing 6M urea and heated prior to SDS-PAGE and Western blotting.

### Western Blotting

For Western blotting of proSAAS, 16% Novex Tris-Glycine gels (Invitrogen) were used for electrophoresis; 10% of each well was applied to the gel. Gels were then electroblotted using the Bio Rad Trans-Blot Turbo RTA Transfer Kit (Bio Rad; #1704271) onto 0.2 μm nitrocellulose membranes at 25V and 1.3A for 15 min. Following transfer, residual proteins in the transferred gel were incubated in Coomassie stain (25% methanol, 10% acetic acid, 0.0003% Brilliant Blue) to validate consistent protein loading. All Coomassie-stained gels and uncropped Western blot images are included in the Supplemental Figures. Prior to proSAAS antiserum addition, nitrocellulose membranes were rapidly rinsed in PBS and subjected to crosslinking in 1% glutaraldehyde (Sigma-Aldrich; Cat #: G5882) in PBS for 15 min with rotation. This cross-linking step has been found to both greatly increase proSAAS immunoreactivity and reduce background (9). Membranes were then rinsed 3 times with PBS to ensure the complete removal of glutaraldehyde and blocked for 1 h in blocking buffer (5% milk (Bio-Rad #1706404) in Tris-Buffered Saline (TBS) containing 0.05% Tween-20 (TBS-Tween). The proSAAS primary antibody consisted of purified IgG from LS46 antiserum, raised in rabbits against recombinant His-tagged 21 kDa proSAAS (9,10,12), and used here at a final concentration of 5 μg/ml.

Western blotting of 7B2, HSP90, and HSP70 was conducted using the same transfer conditions as proSAAS electroblotting; however, glutaraldehyde fixation was not performed. These blots were blocked in 5% milk (Bio-Rad #1706404) in TBS-Tween and the following primary antisera were used: rabbit polyclonal antibody to HSP90 (Cell Signaling; Cat #: 4874S); rabbit polyclonal antibody to HSP70 (Sino Biological; China; Cat #: 103587); and rabbit polyclonal antibody to 7B2 (17), all at a 1:1000 dilution.

For blotting of carboxypeptidase E (CPE), 4-20% Tris-Glycine gels (Invitrogen) were used and electroblotting carried out at 25V and 1.3A for 20 min. Blocking buffer was composed of 2% bovine serum albumin (BSA) (Sigma-Aldrich; Burlington, MA; Cat #: A7906) in TBS-Tween; and goat polyclonal antibody to CPE (R&D Systems; Minneapolis, MN; Cat #: AF3587) was used at a 1:2000 dilution.

All incubations with primary antisera took place overnight at 4°C with gentle rocking. The following day, membranes were washed 3 x with TBS-Tween followed by a 2 h incubation in horseradish peroxidase (HRP)-conjugated secondary antiserum (1:5000 dilution in blocking buffer). Secondary antisera included HRP-linked anti-rabbit antiserum (Jackson ImmunoResearch; West Grove, PA; RRID: AB_2307391) and HRP-linked anti-goat antiserum (Sigma; Cat #: A5420) (CPE only).

Following secondary antiserum washes, membranes were developed using Clarity Western ECL Substrate (Bio Rad #1705061) and imaged using a BioRad cooled camera ChemiDoc Imager. Quantitation of bands was performed using the Bio-Rad imager program Image Lab.

### RNA Isolation, Reverse Transcription, and RT-qPCR

Total RNA was extracted from 4 replicate wells of hippocampal cells in a 12-well plate using Trizol, followed by phase separation using chloroform. The aqueous layer of the phase separation was applied to the column of the Direct-Zol RNA Mini Kit (Zymo; Irvine, CA; Cat #: ZR2070), and isolated following manufacturer’s protocol. RNA concentrations were determined using a Nanodrop spectrophotometer, and 0.5 μg total RNA was reverse transcribed into cDNA using the iScript cDNA Synthesis Kit (Bio-Rad; #1708891). Real-time quantitative polymerase chain reactions (RT-qPCR) were performed in 10 μl reactions containing Power SYBER Green PCR master mix (Bio-Rad; Cat #: 172-5121), 5 ng of cDNA, and 500 nM of the specific forward and reverse primers. RT-qPCR was performed on a CFX96 RT-qPCR system (Bio-Rad) using the following conditions: a 3 min initial denaturation at 95°C, followed by 39 cycles of 15s denaturation at 95°C, 15s annealing at 60°C, and 30s extension at 72°C. Primers used during this experiment included: GAPDH: forward: 5’-CGTGTTCCTACCCCCAAT-3’ reverse: 5’- TGTCATCATACTTGGCAGGTTTCT-3’; proSAAS: forward: 5’-GTC GGC ATT TTG GTG CTG-3’ reverse: 5’-ATT GAG GGC TCA GGG GAT-3’; and CPE: forward: 5’-TGA GAA AGA AGG CGG TCC TAA C-3’ reverse 5’-GCA GAT TGG CAG AAA GCA CAA-3’.

### Immunocytochemistry of Hippocampal Cultures

For visualization of proSAAS within hippocampal cultures, 50,000-100,000 cells were plated on poly-l- lysine-coated coverslips and differentiated for 10 days, with feeding twice a week (9). Cells were rinsed with dPBS, then fixed for 20 min in 4% paraformaldehyde (PFA) at room temperature. After removal of PFA, cells were rinsed 3 times in dPBS, then incubated for 30 min in permeabilization buffer (5% FBS, 0.5% Triton X-100, and 100 mM glycine in dPBS) at room temperature. Cells were then incubated overnight in primary antisera at 4°C in permeabilization buffer. Primary antisera used in this experiment were LS46 (5 μg/ml purified IgG) for visualizing endogenous proSAAS; a mouse monoclonal antibody to Bassoon (UC Davis/NIH; California, USA; Cat #: 75-491); and Acti-Stain^TM^ 670 Phalloidin to mark neurites (200 nM; Cytoskeleton Inc.; Denver, CO; Cat #: PHDN1-A). The following day, cells were rinsed with dPBS prior to a 1 h incubation at room temperature in Alexa-Fluor^TM^ 568 goat anti-rabbit IgG (1:1000; Invitrogen; RRID: AB_10563566) and Alexa-Fluor^TM^ 488 goat anti-mouse IgG (1:1000; Invitrogen; RRID: AB_2534088) in permeabilization buffer. After additional washing with PBS, coverslips were mounted using in-house fluorescent mounting medium to preserve color. To determine the degree of non-specific antibody binding, no-primary control coverslips were treated alongside coverslips used for quantitation, with all other conditions (microscope gain and laser power settings as well as secondary antibody treatment) held constant.

### Analysis of Confocal Images

Coverslips were imaged on a Leica STP 8000 confocal microscope at the magnifications indicated in the figure legends. Gain and laser power settings were determined for each experiment and all images were taken with the same settings and within 24 h of each other. Six Z-stacks of 3 different images were taken per cover slip from random fields; Z stacks were combined into max projections prior to analysis. Quantitation of images was performed blinded using randomized slides. Images containing evident damage or major out of focus regions were eliminated from the analysis, though no more than two images out of six were removed from any one replicate. Analyses were performed using Fiji v1.54 with MorphoLibJ v1.6.2 (18). Unless otherwise stated, default program settings were used during the analysis. To account for background fluorescence, corrected total fluorescence (CTF) for each region of interest (ROI) was calculated by taking the integrated density measurement and subtracting the area selected times the mean background fluorescence [CTF = Integrated density – (area selected x mean background fluorescence)]. For each image, six regions of background from different quadrants were sampled and averaged together to determine the mean background. To analyze the total cellular content of proSAAS immunoreactivity (proSAAS-ir), complete ROIs of actin staining were created via morphological filtering, first by conversion to a binary mask, followed by greyscale closing to remove noise. Finally, morphological reconstruction, with origins centered at the nucleus and with a connectivity setting of 8, was performed to connect discontinuous segments in the ROI using MorphoLibJ. To analyze the soma, nuclei were manually selected using DAPI staining, the resulting ROI was expanded by 3 microns, and proSAAS-ir was quantified. To quantitate the contribution of neurites to total proSAAS-ir, the soma ROI previously mentioned was subtracted from the total cellular ROI. To quantitate synaptic proSAAS-ir, Bassoon immunoreactivity (Bassoon-ir) was thresholded using the “maximum entropy thresholding” algorithm to select only the most intense pixels; and somatic regions were removed to exclude ER- or Golgi-localized Bassoon. Significance was determined by one-way ANOVA with Tukey’s post-test.

### Animals

Two-month-old female 5xFAD mice were obtained from Jackson Laboratories (Bar Harbor, Maine). Prior to surgery, all mice were anesthetized with ketamine (80-150 mg/kg)/xylazine(8-15 mg/kg) via intraperitoneal injection. Mice were monitored for depth of anesthesia by evaluating breathing and response to deep toe pinch. When non-responsive, mice received pain medication (carprofen, 5mg/kg) via subcutaneous injection. Bilateral stereotactic injection of AAVs encoding either proSAAS or eGFP were injected into anesthetized mice into the CA1 region (-1.7 A/P, ±1.3 M/L, and -1.2 D/V; 2.8 x 10^9^ gc/injection site; 0.5 μl/injection site). All mice received subcutaneous injections of carprofen (4-5mg/kg) once daily for 3 days following surgery, as well as topical Lidocaine to the wound twice daily for 3 days following surgery. All mice (10 total: 5 in control group, 5 in experimental group) were then aged for 6 months with weekly weight checks. At 8 months of age, mice were euthanized via intraperitoneal injection of ketamine (500mg/kg)/xylazine (50mg/kg), and immediate transcardiac perfusion with saline followed by 4% paraformaldehyde (PFA) was then performed. Brains were removed and soaked in 30% sucrose at 4 °C for one day prior to flash freezing in Optimal Cutting Temperature compound (OCT; Sakura; Torrance, CA; Cat #: 4583). After freezing, brains were sectioned into 40 μm coronal sections using a Leica cryostat and stored in long-term protectant. All procedures involving animals were approved by the UMB institutional IACUC Committee (Protocol number: 1021010).

All animals remained healthy throughout the duration of the experiment; however, one animal in the eGFP-injected control group was excluded due to an abnormally large plaque number (two-fold greater than all other mice, possibly due to a larger transgene dosage).

### Hippocampal ProSAAS Western Blotting

Hippocampi from 59–61-week-old wild-type and 5xFAD mice were removed on ice, then homogenized in 1 ml of ice-cold extraction buffer (20 mM Tris-HCl, pH 8.0, 100 mM NaCl, 0.5 mM EDTA and 0.5% NP- 40, supplemented with phosphatase inhibitors (Roche; Basel, Switzerland; Cat#: 04906845001) and protease inhibitors (ThermoFisher Scientific; Waltman, MA; Cat# A32955). Homogenization was followed by brief (10 sec) sonication to ensure complete tissue disruption. Samples were then centrifuged at 14000 x g for 5 min at 4 °C. The clear supernatant was removed, a sample was taken for protein determination by the BCA method, and an aliquot of the remaining supernatant was diluted into Laemmli sample buffer containing 6 M urea. Eighty μg of total soluble protein per mouse were loaded onto 16% Novex Tris-Glycine gels (Invitrogen) and Western blotting for proSAAS was performed as described above.

### Immunofluorescence Microscopy

Free-floating sections obtained from PFA-perfused brains of 8- or 12-month-old wild-type and 5xFAD mice were first blocked for 1 h at room temperature in blocking solution (1% BSA, 1% normal donkey serum (Jackson Immuno Research; RRID: AB_2336990) in TBS containing 0.3% Triton-X 100; “TBS-Triton”). All washes took place with gentle rotation. Sections were incubated in Amylo-Glo solution (Biosensis; Australia; Cat #:TR-300-AG) for 20 minutes, followed by a 5 min wash in 0.9% NaCl. Sections were then incubated in primary antiserum [(LS46 IgG (2 μg/ml); Iba1 (ionized calcium binding adaptor molecule) (1:500; Novus; Cat#: 1028SS); glial fibrillary acidic protein (GFAP) (1:500; Millipore Sigma; Burlington, MA; Cat#: 345860)] in blocking solution overnight at 4°C. The following day, sections were washed in TBS-Triton, then incubated in secondary antiserum diluted in TBS-Triton. Secondary antisera included Alexa-Fluor^TM^ 488 donkey anti-rabbit IgG (1:500; Invitrogen; RRID: AB_2535792), Alexa- Fluor^TM^ 555 donkey anti-rabbit IgG (1:500; Invitrogen; RRID: AB_162543), Alexa-Fluor^TM^ 555 donkey anti-goat IgG (1:500; Invitrogen; RRID: AB_2535853), and Alexa-Fluor^TM^ 488 donkey anti-rat (1:500; Invitrogen RRID: AB_2535794). After washing, sections were mounted on gelatin-coated slides and dehydrated in an ethanol series prior to coverslipping with DPX mounting media (Sigma; Cat #: 06522). Sections were imaged on a Nikon Nie fluorescence microscope or a Leica STP 8000 confocal microscope. To determine the degree of non-specific antibody binding, no-primary control slices were treated alongside coverslips used for quantitation, with all other conditions and microscope settings kept constant.

### Plaque Quantitation

For each mouse, three slices surrounding the injection site were imaged at 10x magnification on a Nikon Nie Fluorescent Microscope and analyzed using Nikon NIS-Elements software. Each hippocampus was treated as an independent region of interest (ROI), and each CA1 region was also treated as an independent ROI. Plaque counting and size analysis was performed blinded on randomized slides and Look Up Tables (LUTs) between each image were set identically (50 min, 850 max). Only objects larger than 5 pixels (2.37 μm^2^) and less than 1500 μm^2^ were included in the analysis to reduce artifacts. Plaque burden was calculated as % plaque area / % ROI area. Since 3 slices were used per mouse, and each (independently injected) hippocampus was treated as an independent ROI, we obtained an n=30 for the proSAAS group and n=24 for the eGFP control group (due to the exclusion of one mouse, as described above).

### Statistical Analyses

All statistical analyses were performed using GraphPad Prism 5. Multiple comparisons between groups were performed as described in the figure legends, including unpaired t-test, one-way ANOVA, two-way ANOVA, and Tukey post-tests.

## Results

### ProSAAS is a highly dynamically altered synaptic component following hippocampal homeostatic scaling

To investigate whether proSAAS levels change disproportionately (vs other synaptic proteins) during homeostatic scaling, we treated hippocampal cultures with the sodium channel blocker tetrodotoxin (TTX) to mimic homeostatic upregulation, and with the GABA receptor antagonist bicuculline (BIC) to simulate homeostatic downregulation. Whole cell lysates were Western blotted for immunoreactive proSAAS, carboxypeptidase E (CPE), and 7B2 (**Figure 1A**), and the immunoreactive bands for each were quantitated. ProSAAS was more strongly upregulated upon treatment with TTX (2 µM) than were two other known synaptic vesicle components, namely CPE and 7B2. Across four biological replicates, TTX treatment resulted in a 144% ± 15.4% increase in proSAAS (mean ± SD, n =4 independent experiments, each performed in quadruplicate); a 115% ± 12% increase in CPE; and a 94.8% ± 10.6% change in 7B2 levels (**Figure 1B**). ProSAAS was also more strongly downregulated than these other synaptic proteins upon treatment with BIC. Following a 24 h treatment with BIC (20 µM), proSAAS levels decreased to 38.2% ± 8.6 of the vehicle group, while CPE and 7B2 levels were not significantly altered (CPE levels were 99.1% ± 17.6% that of the vehicle group, while 7B2 was 96.3% ± 7.1%) (**Figure 1B**). Of the three proteins analyzed, proSAAS was the only protein with significant changes following both TTX and BIC treatment (***p<0.0001). Collectively, these data support the idea that proSAAS is more dynamically regulated during homeostatic scaling than are other vesicular components.

**Figure 1.**
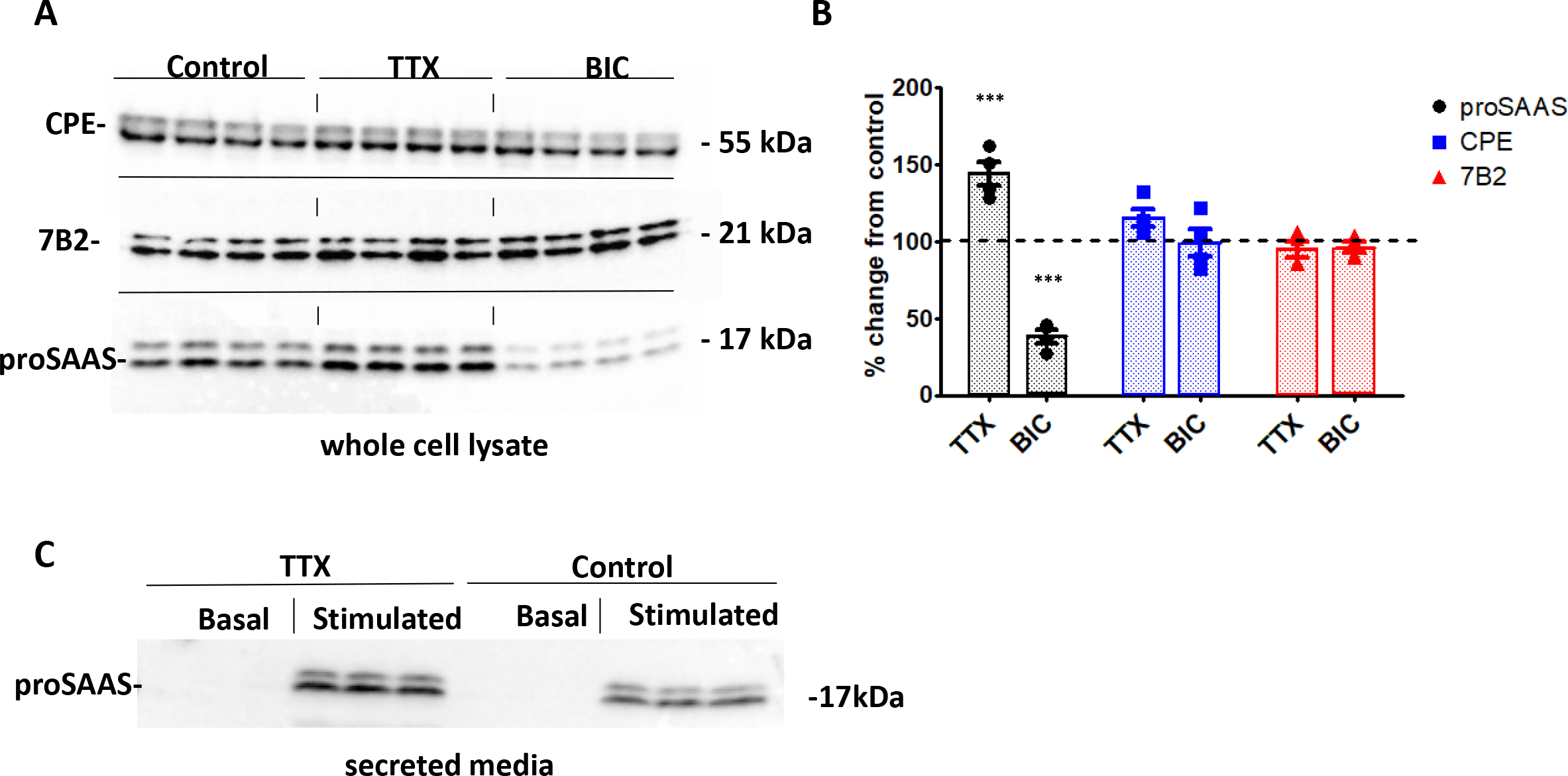
Homeostatic upscaling and downscaling strongly impact proSAAS levels, while other synaptic components remain unchanged. (A) Western blotting of whole cell lysates for synaptic components indicates a rise in cellular proSAAS levels following a 24 h treatment with TTX (2 μM) and decreased proSAAS levels following 24h treatment with BIC (20 μM). Two other synaptic components, CPE and 7B2, remain unchanged. **(B)** Quantitation of Western blots represents the percent change in protein level relative to the TTX vehicle control. Each data point represents an independent experiment; each experiment was conducted in quadruplicate, +/- SD. ***p<0.0001; one-way ANOVA with the Tukey *post-hoc* test when compared to control **(C)** Western blotting of secretion media containing 45 mM KCl confirms that the 17 kDa fragment of proSAAS (LARALL) is released upon stimulation.

To confirm our hypothesis that proSAAS is secreted by differentiated primary hippocampal neurons, we performed Western blotting of the secreted media of stimulated hippocampal neurons for proSAAS. Since primary hippocampal cultures possess a tightly regulated secretory pathway, neurons were subjected to depolarization using KCl in order to obtain detectable amounts of secreted proSAAS in the medium. Upon stimulation with 45 mM KCl, the 17 kDa fragment of proSAAS (which we term “LARALL” after an amino acid sequence contained within it) was secreted into the medium. As expected, when neurons were treated with TTX (2 µM) for 24 h prior to stimulation, proSAAS levels increased in the secretion medium (**Figure 1C)**. These data show that the 17 kDa LARALL fragment is the primary immunoreactive form of proSAAS produced and secreted by these hippocampal cultures. The LS46 antiserum, raised against recombinant 21 kDa proSAAS, is known to detect both the 21 kDa and the 27 kDa (intact) forms of proSAAS (9), yet these larger species were found neither in cells nor in secretion media, suggesting complete processing of proSAAS to this internal fragment.

### GABA agonists exert changes similar to tetrodotoxin; heat shock protein levels remain unchanged

We then treated hippocampal cultures with the GABA agonist muscimol for 24 h to determine if this would result in effects opposite to those produced by the GABA antagonist BIC, which strongly reduced proSAAS content. Using Western blotting and band quantitation, we saw a significant increase in proSAAS levels using both 5 µM (p<0.05) and 10 µM (p<0.001) muscimol treatments (**Figure 2A-B**).

**Figure 2.**
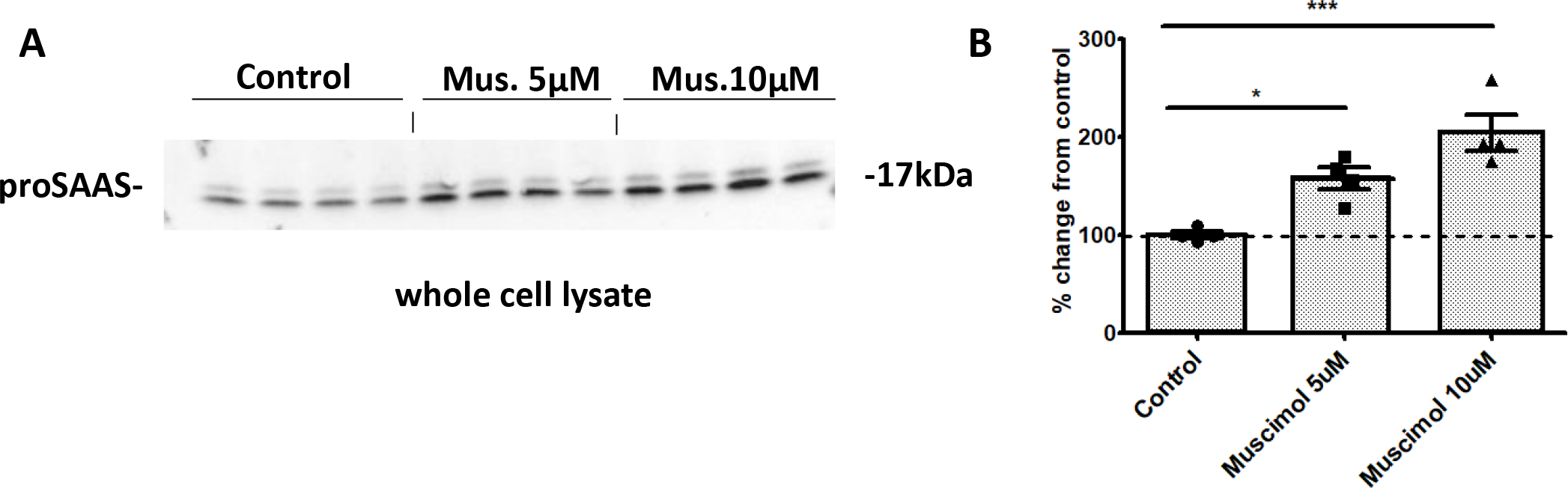
GABA agonists are as potent as TTX in increasing proSAAS levels. (A) Hippocampal cultures were treated with 5 μM or 10 μM muscimol for 24 h and whole cell lysates were then Western-blotted for proSAAS. **(B)** Quantitation indicates a significant increase in proSAAS levels at both 5 μM and 10 μM muscimol concentrations. Data are presented as the mean +/- SD of 4 replicates. *p<0.05, ***p<0.0001; one-way ANOVA with the Tukey *post-hoc* test.

The synaptic specificity of these homeostatic scaling changes was tested by examining the levels of two cytoplasmically-expressed heat shock proteins, HSP70 and HSP90, using a similar Western blotting procedure. Treatment of hippocampal cultures with TTX, BIC (or, in a separate experiment, muscimol) generated no significant changes in the levels of these two heat shock proteins (**Figure 3 A, B**).

**Figure 3.**
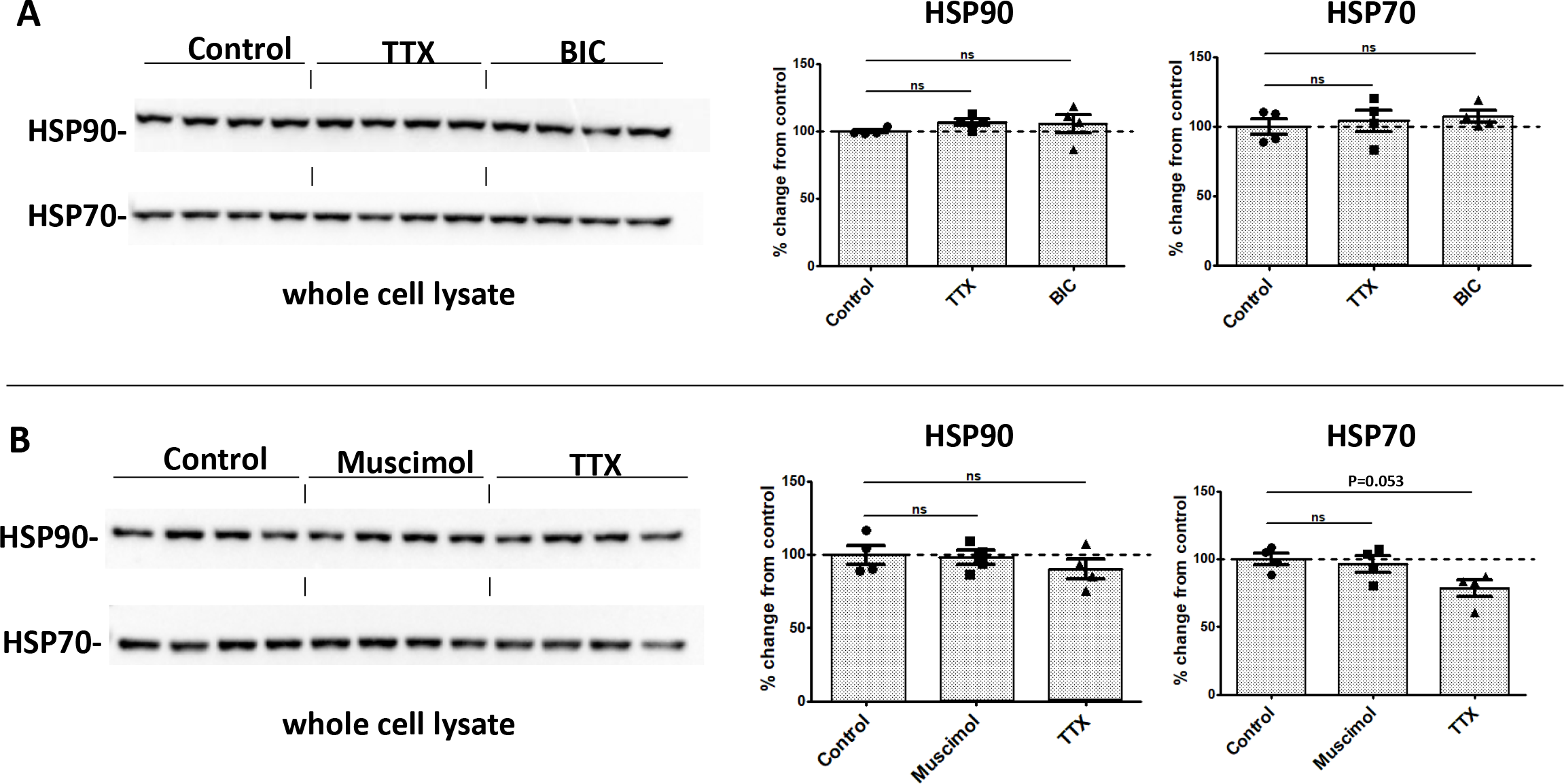
Cytoplasmic heat shock proteins levels are not altered by treatment with TTX or GABA related-drugs. (A) Following treatment with TTX (2 μM), BIC (20 μM), HSP90 and HSP70 levels in whole cell lysates remain constant. **(B)** An independent experiment employing muscimol (10 μM) yielded similar results. Data are presented as the mean +/- SD of 4 replicates; one-way ANOVA with the Tukey *post-hoc* test. ns, not significant.

### ProSAAS protein changes are larger than proSAAS mRNA changes, suggesting possible translational regulation or inhibition of degradation

To determine if the increase in proSAAS levels upon TTX treatment is due to increased mRNA production, increased mRNA usage, or both, we conducted a protein and mRNA time course analysis using TTX-treated hippocampal cultures. At various time points, we performed either Western blotting or qPCR on samples plated in parallel. **Figure 4A** shows dynamic proSAAS protein increases at 24, 48 and 72 h, while mRNA levels remained largely static. We saw an increase in proSAAS protein at 48 h after initial TTX treatment (166 ± 28 %) and the greatest increase 72 h after treatment (171 ± 32%). By contrast, over the course of these same 72 h, CPE protein levels and mRNA levels both remained largely static (**Figure 4B**).

**Figure 4.**
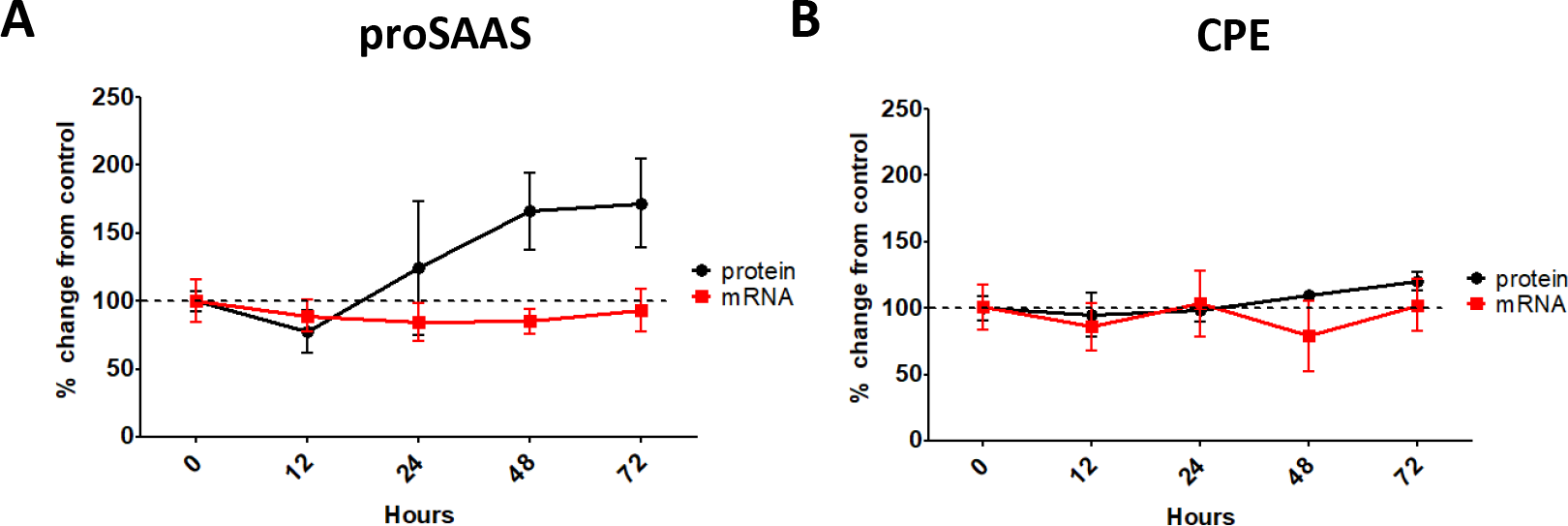
Time course of homeostatic scaling indicates dynamic proSAAS protein changes, while mRNA levels remain unchanged. Differentiated hippocampal cells were treated with 2 μM TTX and harvested at the time points indicated. Protein levels were determined by Western blotting of whole cell lysates; and mRNA levels were determined by quantitative real-time PCR. Panel **(A)** illustrates dynamic proSAAS protein changes and static mRNA levels, while panel **(B)** shows static CPE protein and mRNA levels over the course of 72 hours. Each mRNA species is shown relative to GAPDH and is plotted as the fold-increase compared to the vehicle control. The data represent the average of 4 biological replicates, each containing 3 technical replicates, +/- SD.

### ProSAAS homeostatic scaling occurs preferentially in the synaptic area

To investigate the distribution of proSAAS upregulation during homeostatic scaling, primary hippocampal cultures were plated on coverslips and treated with TTX (1 µM). Since hippocampal cultures were more sparsely plated for morphometric analysis, a decreased TTX concentration was used in these experiments to preserve neuronal health while still stimulating upregulation. Fixed cultures were treated with antibodies against proSAAS, the presynaptic marker protein Bassoon, DAPI nuclear staining, and the actin stain phalloidin to mark neurites. **Figure 5A** shows example images of hippocampal cultures 12 and 24 h post TTX treatment stained for proSAAS *(green*) and Bassoon (*red*). Different plating densities between the immunocytochemical and Western blotting experiments, along with differences in the TTX concentrations used for the two types of experiments, may have contributed to variations in proSAAS levels between Western blotting and immunocytochemical experiments.

**Figure 5.**
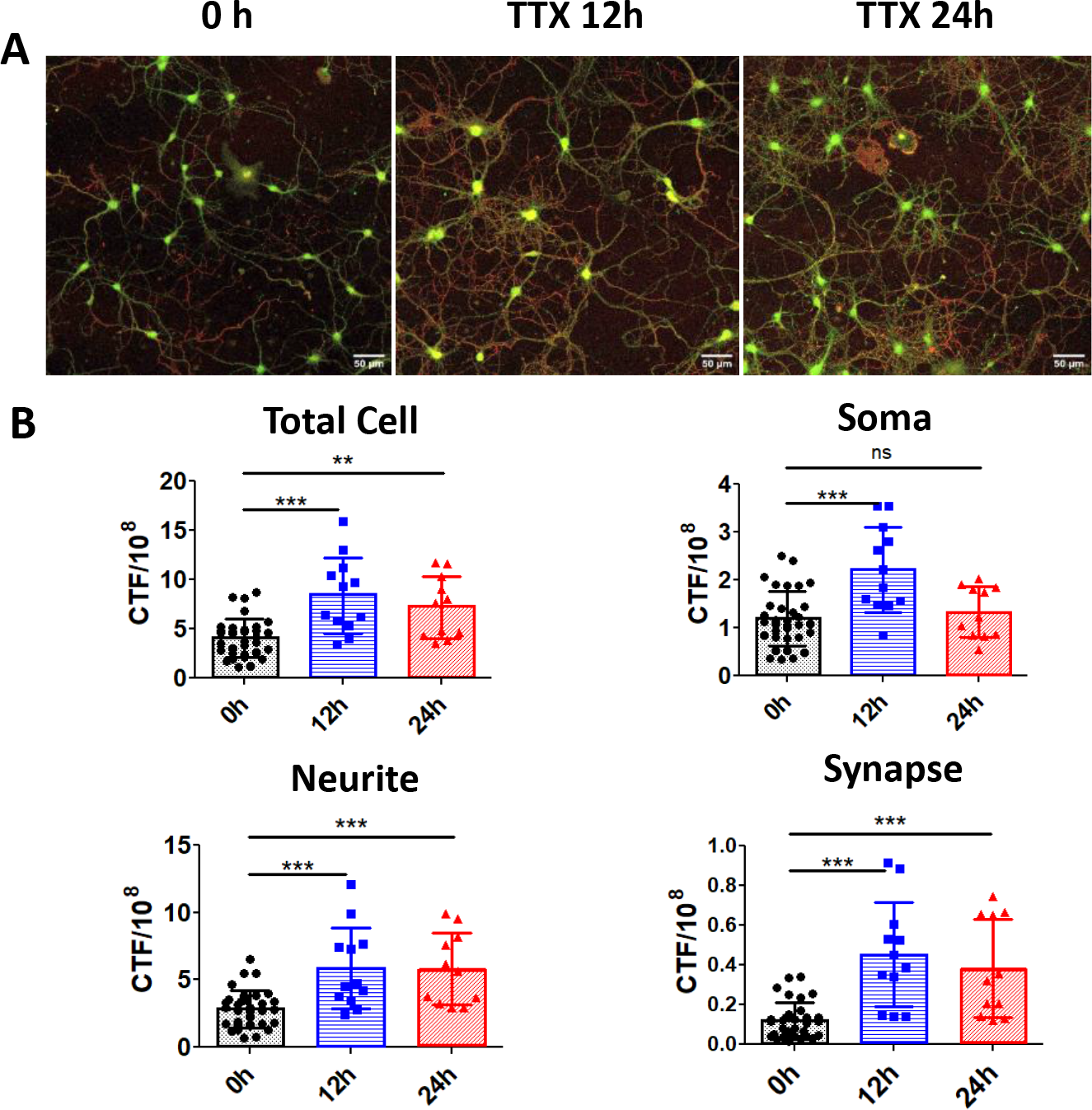
Confocal microscopy of TTX-treated cells indicates that both cell body and neurites exhibit increased proSAAS-ir after TTX. Primary hippocampal cells were cultured for 10 days and treated with TTX (1 μM) or the vehicle control for 12 or 24 h. Each point represents a separate image, with 6 images per coverslip and 2 biological replicates. **(A)** Example images of immunoreactive proSAAS (*green*) and the presynaptic marker Bassoon (*red*), showing increased proSAAS fluorescence following TTX treatment (50 μm scale bars). **(B)** Quantitation of proSAAS fluorescence in different cellular compartments across 0, 12 and 24 h. 0 h: n = 33, 12 h: n = 12, 24 h: n = 11. *p<0.05; **p<0.01; ***p<0.001; one-way ANOVA with the Tukey *post-hoc* test. A no-primary antibody control of hippocampal cultures is shown in Supp. Fig 4. CTF, corrected total fluorescence.

As expected based on Western blotting results, confocal imaging analysis confirmed that proSAAS-ir increased across the total neuronal (actin-stained) area at 12 and 24 h (**Figure 5B**). We found a significant increase in proSAAS-ir in neurites at 12 and 24 h (p <0.0001) as well as in synaptic areas (as judged by proximity to Bassoon) at 12 and 24 h of TTX treatment (p <0.0001). Interestingly, we detected increased proSAAS staining in neuronal soma at 12 h of TTX treatment, but not at 24 h. Taken together with our finding that synaptic proSAAS is increased during this same time period, we surmise that proSAAS translocation from the soma to the synapse may occur during this time.

### Immunoreactive proSAAS is enriched in the hippocampal mossy fiber layer in wild-type mice and is associated with all amyloid plaques in 5xFAD mice

We performed fluorescent staining of coronal sections to visualize the location of proSAAS-ir in normal mouse brain. While, as previously found (8,19,20)(22,23), proSAAS expression is widespread within the brain, we observed particularly rich proSAAS staining in the mossy fiber tract projecting from the dentate gyrus to the CA3 region of the hippocampus, specifically within the hilar region (**Figure 6A**).

**Figure 6.**
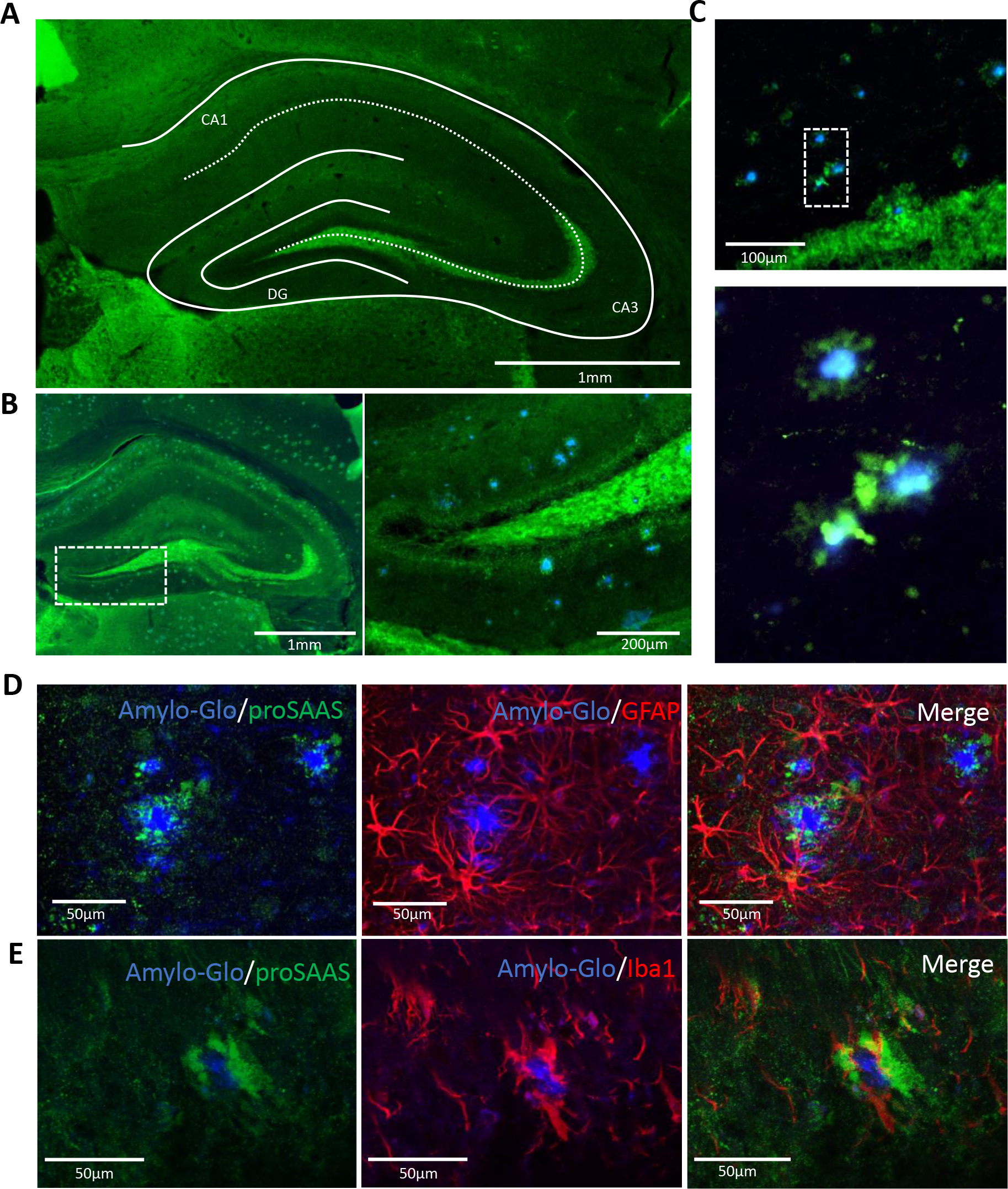
Endogenous proSAAS is enriched in the mossy fiber tract of the hippocampus and surrounds amyloid beta plaques in 5xFAD mice. (A) Coronal section of a WT 12-month-old female mouse brain stained for proSAAS-ir (*green*). **(B)** Coronal section of a 12-month-old female 5xFAD mouse brain stained for proSAAS-ir (*green*) and amyloid plaques using Amylo-Glo (*blue*), and further magnified (**C**). (**D**) Coronal sections of a 12-month-old female 5xFAD mouse show that plaque (*blue*)-associated proSAAS (*green*) does not colocalize with glial cells (*GFAP; red*) nor with microglia (**F**) *(Iba1; red)* (50 μm scale bars). A no-primary antibody control was used to determine nonspecific staining (Supp. Fig. 5).

Pulldown studies clearly show that proSAAS binds amyloid (12,21). We examined the distribution of endogenous proSAAS in 12-month-old 5xFAD mice. ProSAAS-ir is similarly rich in the mossy fiber tract in the 5xFAD mouse hippocampus, but in addition, in 5xFAD mice, immunoreactive proSAAS is present in the vicinity of every amyloid plaque, often as a rosette surrounding the plaque (**Figure 6B**). A magnified image shows this apparent proSAAS sequestration of amyloid (**Figure 6C**).

The great majority of brain proSAAS is synthesized by, and secreted from neurons (8,22,23). To confirm that proSAAS-ir surrounding extracellular plaques likely derives from secreted material, i.e. is not present in plaque-associated cells, we stained adult 5xFAD hippocampal slices for proSAAS, amyloid plaques (Amylo-Glo), and the glial cell marker GFAP. ProSAAS-ir (*green*) clearly does not colocalize with glial cells (*red*) associated with amyloid plaques (*blue*) (**Figure 6D**). Additionally, staining with antiserum to the microglial marker Iba1 (*red*) shows no colocalization of proSAAS with microglia (*green*) (**Figure 6E**), confirming that these cells are also not the source of plaque-associated proSAAS.

Plaque-associated proSAAS-ir was found in two different morphologies, either within rosette-type formations, or within larger globular aggregates. **Figure 6D** shows an example of the rosette morphology, while in **Figure 6E** the more globular morphology is shown. Additional Z-stack imaging of plaques (not shown) revealed that even when proSAAS-ir completely surrounded plaques, it was never present within the Amylo-Glo-stained dense core, suggesting that it is not involved in early events in plaque formation.

### Overexpression of proSAAS in the CA1 of the hippocampus results in decreased amyloid plaque size and plaque burden in 5xFAD mice

The above immunohistochemical demonstration of proSAAS association with plaques in the 5xFAD mouse (**Figure 6),** taken together with previous work showing that proSAAS blocks amyloid aggregation in *in vitro* experiments (12) led to the hypothesis that proSAAS association with plaques might impact plaque growth. To quantify possible anti-aggregation effects of overexpressed proSAAS, 2-month-old 5xFAD female mice received bilateral CA1 hippocampal injections of AAVs encoding either proSAAS or eGFP (control); female 5xFAD mice were used due to their increased plaque size and number compared to males (24). Mice were then aged 6 months to allow sufficient plaque formation (**Figure 7A**). eGFP-encoding virus was used to verify injection specificity to the C A1 region (**Figure 7B**). Virally-mediated proSAAS overexpression in the CA1 region was confirmed 6 months after virus injection at the time of plaque analysis (**Figure 7C**).

**Figure 7.**
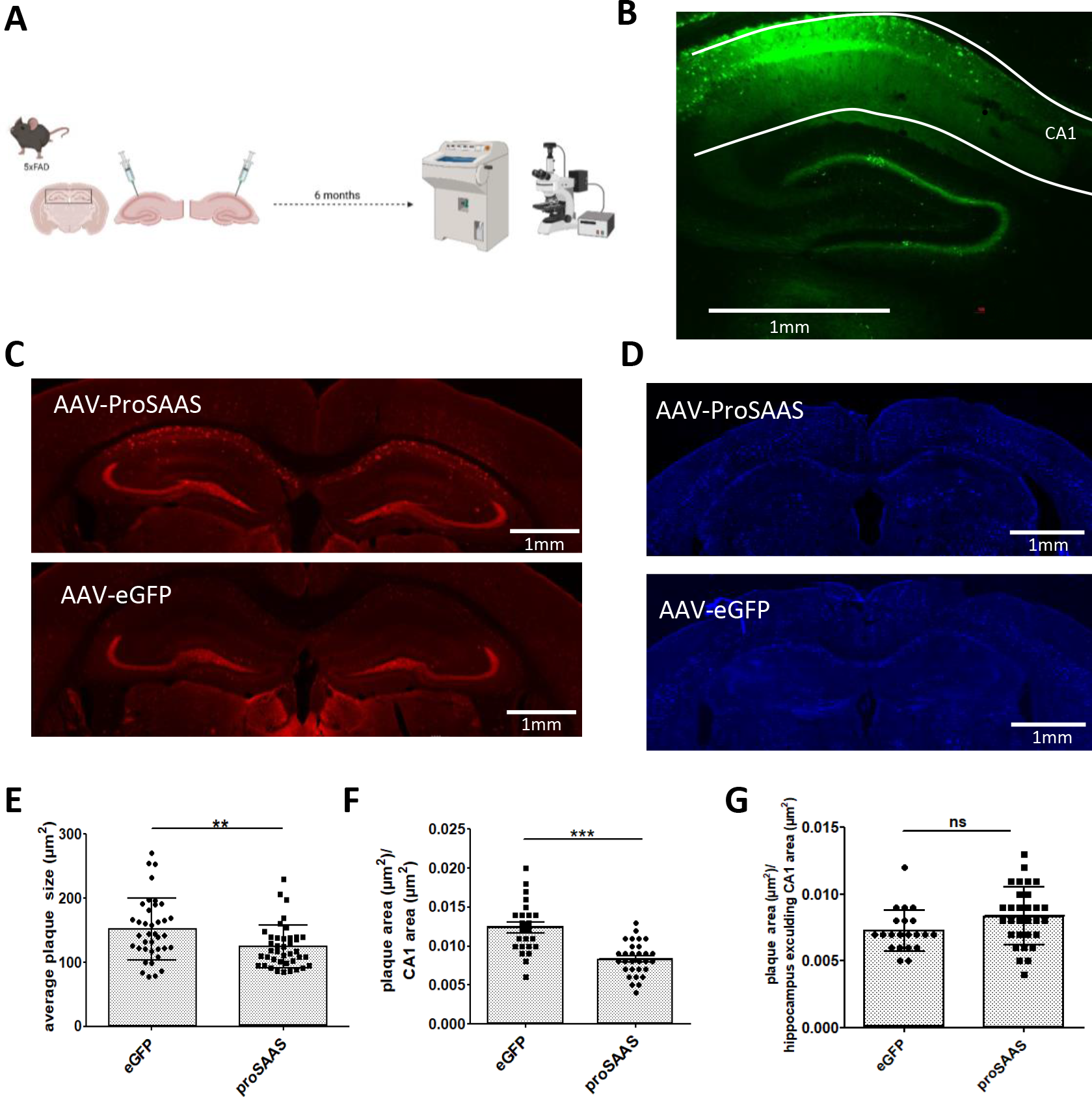
ProSAAS overexpression in the CA1 region results in decreased plaque size. (A) The experimental workflow shown illustrates bilateral stereotactic injections of 5xFAD mice of AAVs encoding either eGFP (control) or proSAAS. Mice were then aged 6 months prior to perfusion, cryosectioning, immunohistochemistry, and plaque analysis. **(B)** A coronal section imaged for GFP confirms proper virus injection to the CA1 region. **(C)** Confirmation of AAV2/1-mediated proSAAS overexpression in the CA1 via proSAAS staining (shown in *red*). (**D**) Example images of coronal sections stained using Amylo-Glo, the stain used for plaque quantitation. **(E)** ProSAAS overexpression results in a significantly decreased average plaque size in the CA1. (**F**) Plaque burden in the hippocampal areas remote from AAV injections resulted in no significant differences between the control and experimental groups. By contrast, plaque burden in the CA1 was significantly decreased by proSAAS overexpression **(G)**; n=30 for proSAAS group, n=24 for eGFP group (see Methods). Each data point represents the average plaque size in a single hippocampus obtained from one hemisphere. **p<0.01, ***p<0.001; unpaired t-test.

Forty-micron hippocampal sections surrounding the injection site were then stained for amyloid plaques and size and number were quantitated in each slice. All staining, imaging, and quantitation was performed blinded to experimental condition; representative images of the Amylo-Glo stain used for quantitation are shown in **Figure 7D**. These experiments revealed a significant decrease in plaque size in the CA1 region in proSAAS-overexpressing 5xFAD mice, with the GFP control group averaging 152 ± 8 µm^2^ and the proSAAS-overexpressing group averaging 124 ± 5 µm^2^ (p<0.01) (**Figure 7E**).

Interestingly, analyses of CA1 plaque burden reveal a significantly increased plaque burden in the GFP-expressing control group (0.012 plaque area µm^2^/CA1 µm^2^) as compared to the proSAAS-overexpressing group (0.008 plaque area µm^2^/CA1 µm^2^) (**Figure 7F**). We also examined plaque burden in areas remote from viral proSAAS overexpression by calculating this parameter in the entire hippocampus and then excluding plaques within the CA1 region. Using this measurement, we found no significant difference in plaque burden between the eGFP-expressing group and the proSAAS-expressing group (**Figure 7G**).

## Discussion

Compensatory synaptic adaptation in response to changes in neuronal activity is thought to maintain synaptic efficiency by increasing the levels and locations of proteins required for neurotransmission following upregulation of network activity, and decreasing these same parameters when activity is low. Modeling of this “homeostatic response” has been performed in hippocampal cultures using the sodium channel blocker TTX, which effects a reduction in network activity, as well as with the GABAergic inhibitor bicuculline, which acts to restrain inhibitory GABAergic input - thus causing network hyperactivity. Homeostatic upregulation and/or redistribution of proteins in response to TTX has been demonstrated both in specific neuronal circuits (25) as well as in proteomics studies of hippocampal cultures (7). Perhaps not surprisingly, many of the largest changes in TTX-upregulated proteins occur in synaptic vesicle components, for example Scg3 (secretogranin 3), 7B2 (also known as secretogranin 5), and CPE (7). Interestingly, in these proteomics studies, the synaptic vesicle protein that consistently exhibited the highest increase after TTX treatment was the proSAAS chaperone (7,26). The fact that this particular protein, whose synaptic functions are as yet undefined, should be so highly regulated during homeostatic scaling is intriguing.

To obtain information on proSAAS changes during homeostatic scaling, we first validated the prior findings by performing proSAAS protein and RNA measurements in similar hippocampal primary cultures. Our work confirms that proSAAS indeed exhibits disproportionate homeostatic changes as compared to other synaptic components. However, some discrepancies with the former proteomics studies do exist; for example, the prior proteomics studies indicate much larger fold-increases after TTX treatment (249% *vs* our 144%) (7). Additionally, we did not observe robust upregulation of 7B2 or CPE, which were also top hits in the prior proteomics study (7). Interestingly, we saw a significant decrease in proSAAS upon BIC treatment (to 38% of the vehicle group), while in the prior proteomics study, no significant change in proSAAS levels was detected following BIC. Differences in the sensitivity of protein detection between our Western blotting protein measurements and proteomics methods - which included isotopic protein labeling and mass spectrometric quantitation- may account for some of these discrepancies, though does not explain the differential sensitivity to BIC.

Our qPCR results indicating stable mRNA expression despite dynamic changes in protein also support the prior proteomics study, which found that 24 h of TTX treatment results in both increased proSAAS synthesis and reduced proSAAS degradation (7). Our confocal analysis, which indicates a synaptic proSAAS increase with no somatic increase at 24 h after TTX treatment supports potential proSAAS translocation to neurites/synapse as well as increased synaptic translation of proSAAS (or decreased degradation). The prior proteomics study indicated that both increased synthesis as well as decreased degradation play a role in the increased proSAAS levels seen after homeostatic upscaling (7). If synaptic translation of proSAAS is indeed increased, the question arises as to how full-length synaptically- synthesized proSAAS can be so efficiently processed to the 17 kDa internal segment LARALL, since proteolytic processing is thought to occur through the action of Golgi and secretory granule proprotein convertases (8,27). These results, which may support synaptic precursor processing, warrant further study.

Interestingly, LARALL, which contains a highly conserved sequence (10,28), is the largest form of proSAAS stored by differentiated hippocampal cultures and is readily released by hippocampal neurons upon depolarization. With the knowledge that this form contains the active anti-aggregant portion of proSAAS (12), we hypothesize that LARALL is released into the extracellular space to chaperone new synaptic proteins during periods of intense protein synthesis - such as those occurring during homeostatic scaling.

Homeostatic scaling is necessary to maintain synaptic function within a physiological limit; thus, this scaling capability is of great interest in the context of neurodegenerative diseases such as AD (4). As AD progresses, accumulation of toxic amyloid oligomers in the hippocampus results in altered thresholds for LTP and LTD; synapse loss; and impairment of homeostatic scaling (6,29,30). As mentioned above, proSAAS functions as an amyloid anti-aggregant, both *in vitro* as well as in cell models of amyloid toxicity (12), and has been shown to bind amyloid (12,21). Additionally, at least 6 different proteomics groups have shown that proSAAS CSF levels drop during disease progression [see meta-analysis in (31)], supporting possible brain retention of proSAAS during AD development. Recent transcriptomics studies have found increased levels of proSAAS mRNA in both human AD brain samples (32) as well as in two mouse models of AD (33–35). Interestingly, proSAAS was specifically identified as a neuronal “plaque-induced gene” in the spatial transcriptomics study of Zeng et al. (33). However, we were not able to detect increased levels of proSAAS-derived peptides in soluble extracts of 5xFAD mouse hippocampus using Western blotting (Supplemental Figure 6). The reasons behind this are unclear at present.

Our hippocampal immunohistochemistry data show that the mossy fiber pathway, which carries information from the CA3 region to the dentate gyrus, contains extremely high levels of proSAAS. Others have shown specific localization of proSAAS transcripts in hippocampal excitatory neurons (35). The enhanced presence of proSAAS within the mossy fiber tract is intriguing considering the clear involvement of this pathway in both learning and memory as well as in homeostatic scaling (25). Thus, the enrichment or loss of proSAAS in this specific pathway is likely to be associated with functional consequences. Electrophysiological experiments employing proSAAS-encoding (or reducing) viral reagents will be useful to directly investigate synaptic effects of proSAAS expression or loss during homeostatic scaling.

The reduction in amyloid plaque burden seen with proSAAS overexpression, taken together with the large protein changes during homeostatic scaling, indicate that this small chaperone protein may also act to preserve synaptic function and neuronal health during AD progression, similar to its profoundly protective effects in a Parkinson’s disease model (11). Other secreted proteins, such as the astrocyte-derived chaperone clusterin, have also been shown to play a role in Aβ clearance and excitatory synaptic transmission (36) as well as in other neurodegenerative diseases [reviewed in (37,38)], although, unlike proSAAS, clusterin overexpression is associated with increased (and redistributed) plaque deposition (39). Of note, the synapsin promoter driving proSAAS expression in our hippocampal viral overexpression experiments results only in neuron-specific expression; more robust anti-plaque effects might be obtained if a stronger promoter, for example CMV (cytomegalovirus) had been employed, as in (36). Future experiments should involve the comparison of different promoters to determine whether amyloid-bound proSAAS must necessarily first be secreted from neurons; or whether support cells such as glia might also be artificially recruited to express proSAAS, thus increasing extracellular proSAAS levels. In a best-case scenario, increasing brain proSAAS expression via use of this strong promoter might further limit plaque size and reduce plaque clustering (40).

In summary, the functional consequences of proSAAS-mediated plaque sequestration are still unclear; additional experiments will be required to show that proSAAS overexpression is indeed neuroprotective in the 5xFAD model mouse. However, given the profound protective effects of proSAAS overexpression in a rat Parkinson’s disease model (11), we predict that both hippocampal network homeostasis as well as cognition will be functionally improved by hippocampal proSAAS overexpression in AD mouse models.

## Acknowledgements

We are extremely grateful to Minerva Contreras for her excellent preparation of primary rat hippocampal cells. We thank Shuxin Zhao and Dr. Celine Plachez for their help with cryosectioning and immunohistochemistry; and Dr. Reha Erzurumlu for the use of his Nikon microscope. We also thank Dr. Manita Shakya, Praise Apanisil and Dr. Tripti Joshi for preliminary experiments; and Dr. Ramesh Chandra of the University of Maryland Virus Core Facility for AAV2/1 virus production. This work was supported by NIH grant AG062222 to I.L.

## Authors Contributions

IL and SM wrote and conceptualized the manuscript. SM, NS, CH and SN performed experiments. Animal and immunohistochemical work was done by SM. Immunocytochemistry and confocal quantitative analysis was performed by NS and SM. Editing was done by IL and SM.

## Data Availability Statement

The data that support the findings of this study, as well as all in-house constructed reagents, are available from the corresponding author upon reasonable request.

