## supplemental figures for "ProSAAS is Preferentially Secreted from Neurons During Homeostatic Scaling and Reduces Amyloid Plaque Size in the 5xFAD Mouse Hippocampus"

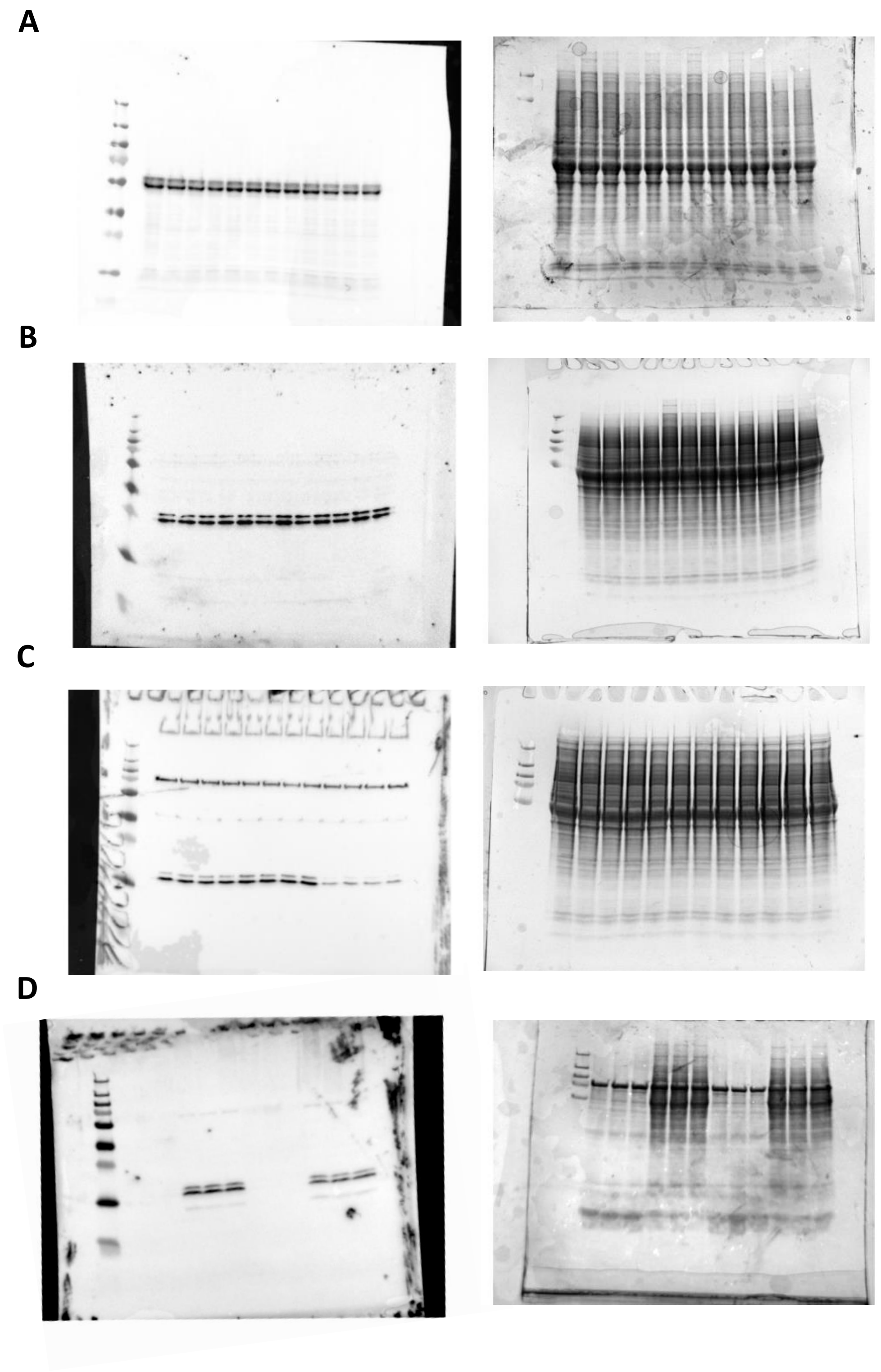


**Supp. Figure 1**. **Uncropped Western blot and corresponding Coomassie-stained gel of TTX and/or BIC treated hippocampal cells in Figure 1**. Residual gel proteins following transfer were stained with Coomassie and imaged (*right*) alongside the uncropped Western blots (*left*) for CPE (**A**), 7B2 (**B**), and proSAAS (**C**), as seen in **Figure 1A**. Panel (**D**) shows the uncropped Western blot (*left*) and residual gel stained with Coomassie (*right*) from the proSAAS secretion experiment presented in **Figure 1C**.

**ProSAAS is Preferentially Secreted from Neurons During Homeostatic Scaling and Reduces Amyloid Plaque Size in the 5xFAD Mouse Hippocampus** Samira Mitias, Nicholas Schaffer, Saaya Nair, Chelsea Hook and Iris Lindberg

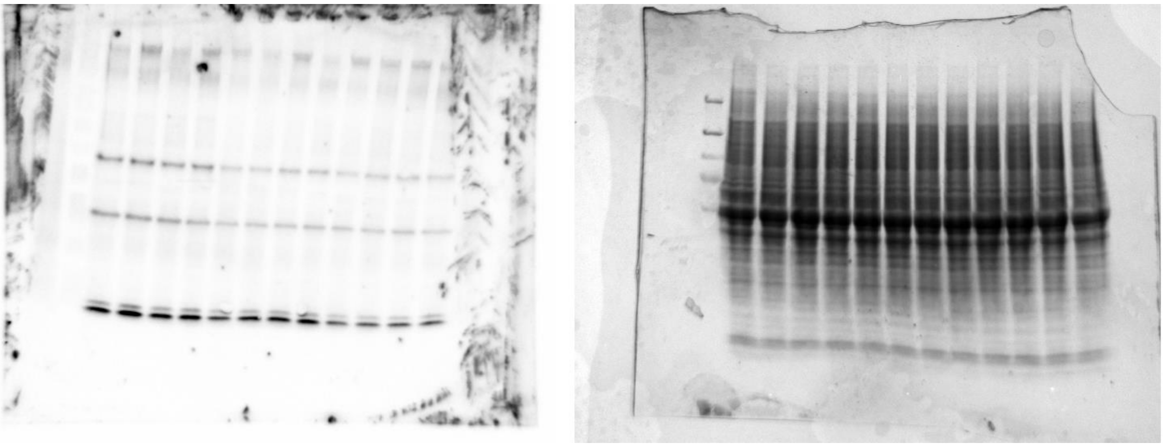

**Supp. Figure 2. Uncropped Western blot and corresponding Coomassie-stained gels of TTX- and muscimol- treated hippocampal cells in Figure 2.** Residual gel proteins were stained with Coomassie and imaged (*right*) alongside the uncropped Western blot (*left*) for proSAAS, as seen in **Figure 2A**.

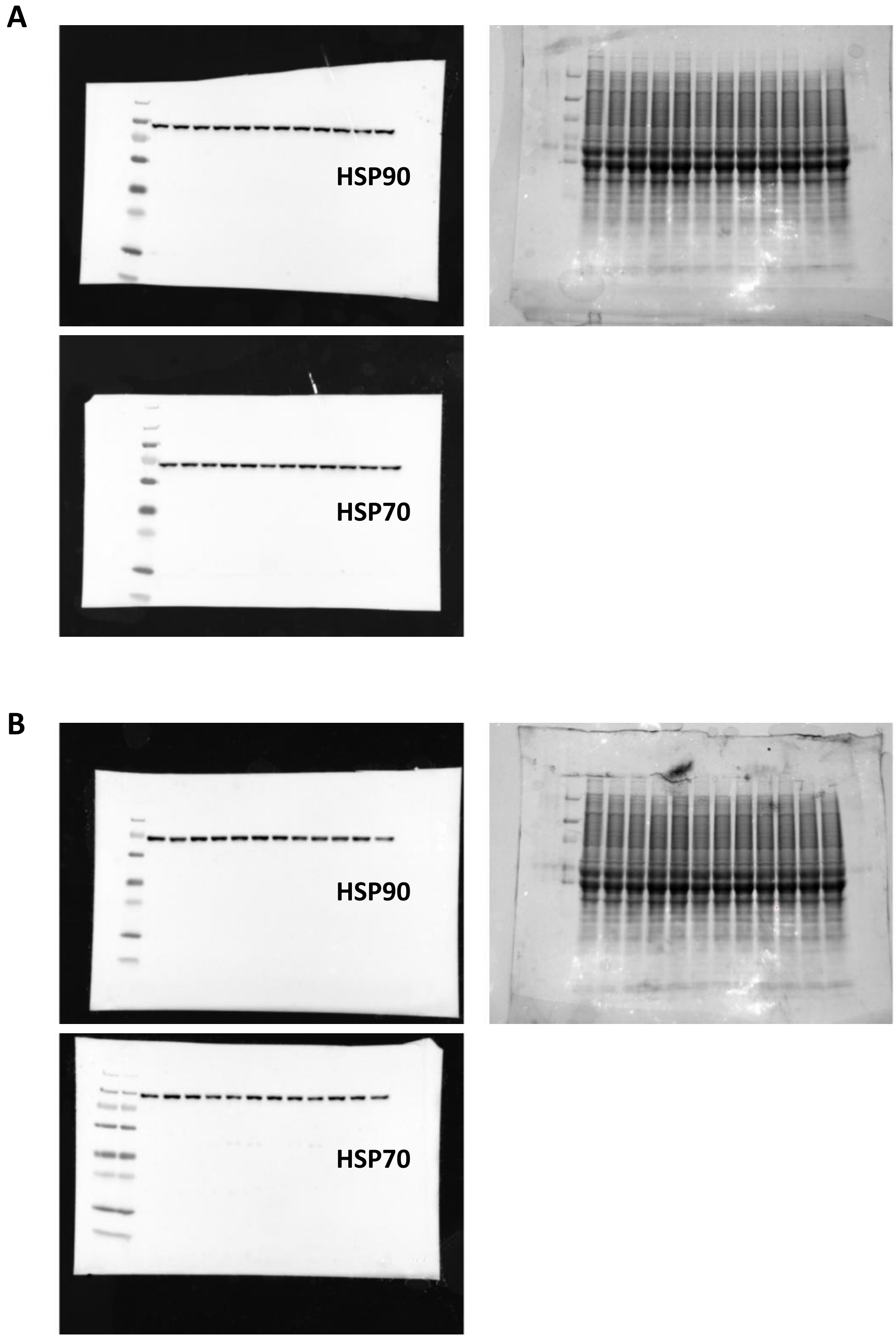


**Supp. Figure 3. Uncropped Western blot and corresponding Coomassie-stained gels of TTX, muscimol, and/or BIC treated hippocampal cells in Figure 3.** Residual proteins were stained with Coomassie and imaged (*right)* alongside the uncropped proSAAS Western blot (*left*) as seen in **Figure 3A** and **3B. Supp**. **Figure 3A** shows the uncropped Western blots and Coomassie gel that generated **Figure 3A**, and **Supp. Figure 3B** shows the uncropped Western blots and Coomassie gel that generated **Figure 3B**.

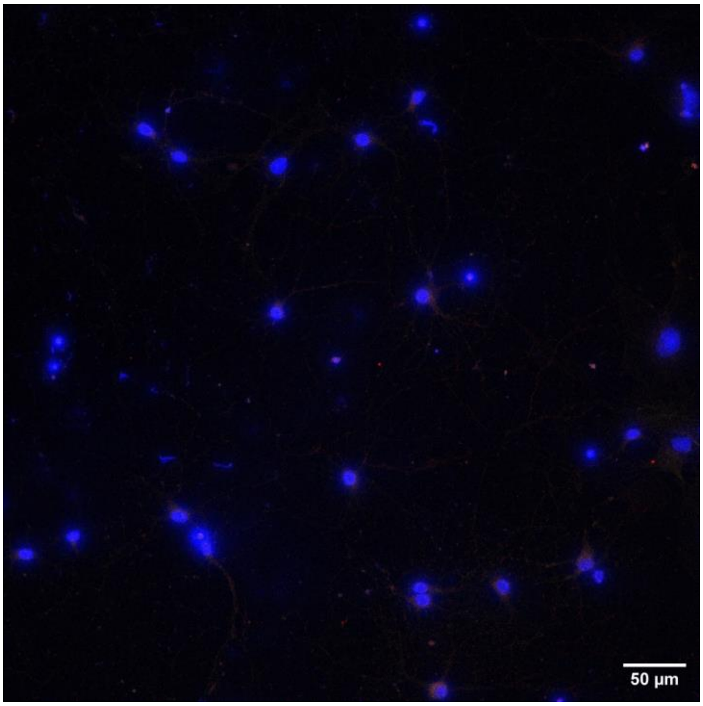


**Supp. Figure 4. No-primary control of confocal hippocampal culture images.** Coverslips of hippocampal cultures grown on glass coverslips and fixed for immunocytochemistry were used to generate **Figure 5**. In parallel with this experimental group, this figure shows control hippocampal cell coverslips which were treated with solutions lacking primary antibody but incubated in secondary antibody and DAPI (to determine the degree of non-specific staining).

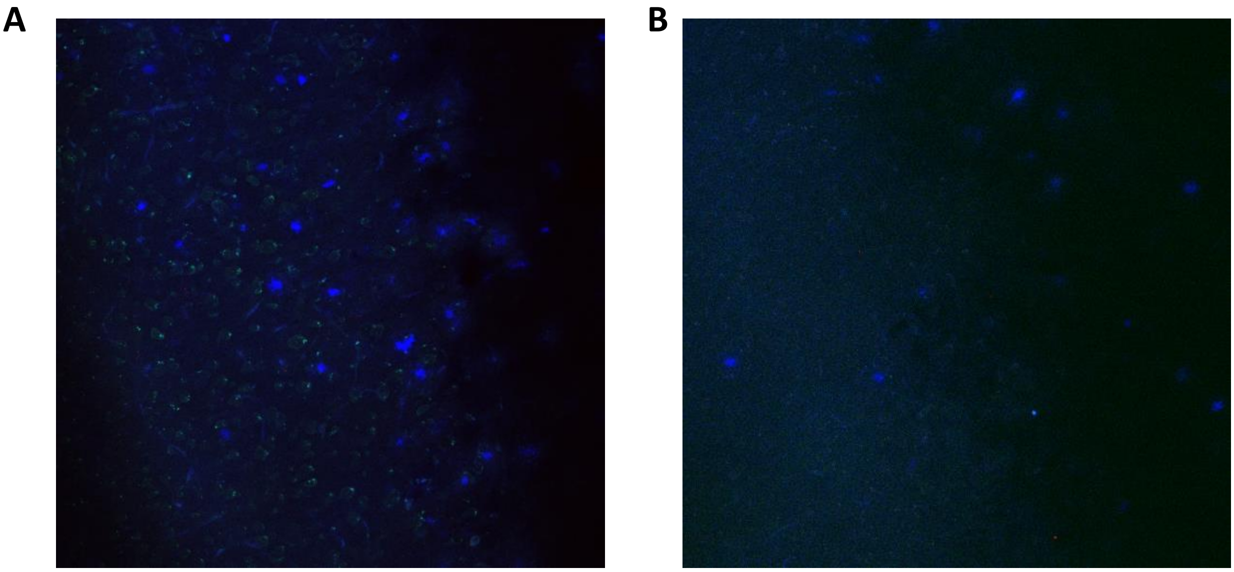

**Supp. Figure 5. No-primary control of hippocampal slices.** (**A**) A coronal section of 5xFAD mouse brain was treated alongside the experimental slice that generated **Figure 6E**, but in the absence of GFAP and proSAAS primary antibodies. (**B**) A similar coronal section was treated alongside the experimental slice that generated **Figure 6E**, but in the absence of Iba1 and proSAAS primary antibodies. All no-primary control slices were imaged at the same time and with the same microscope settings as the images shown in **Figure 6E-F**.

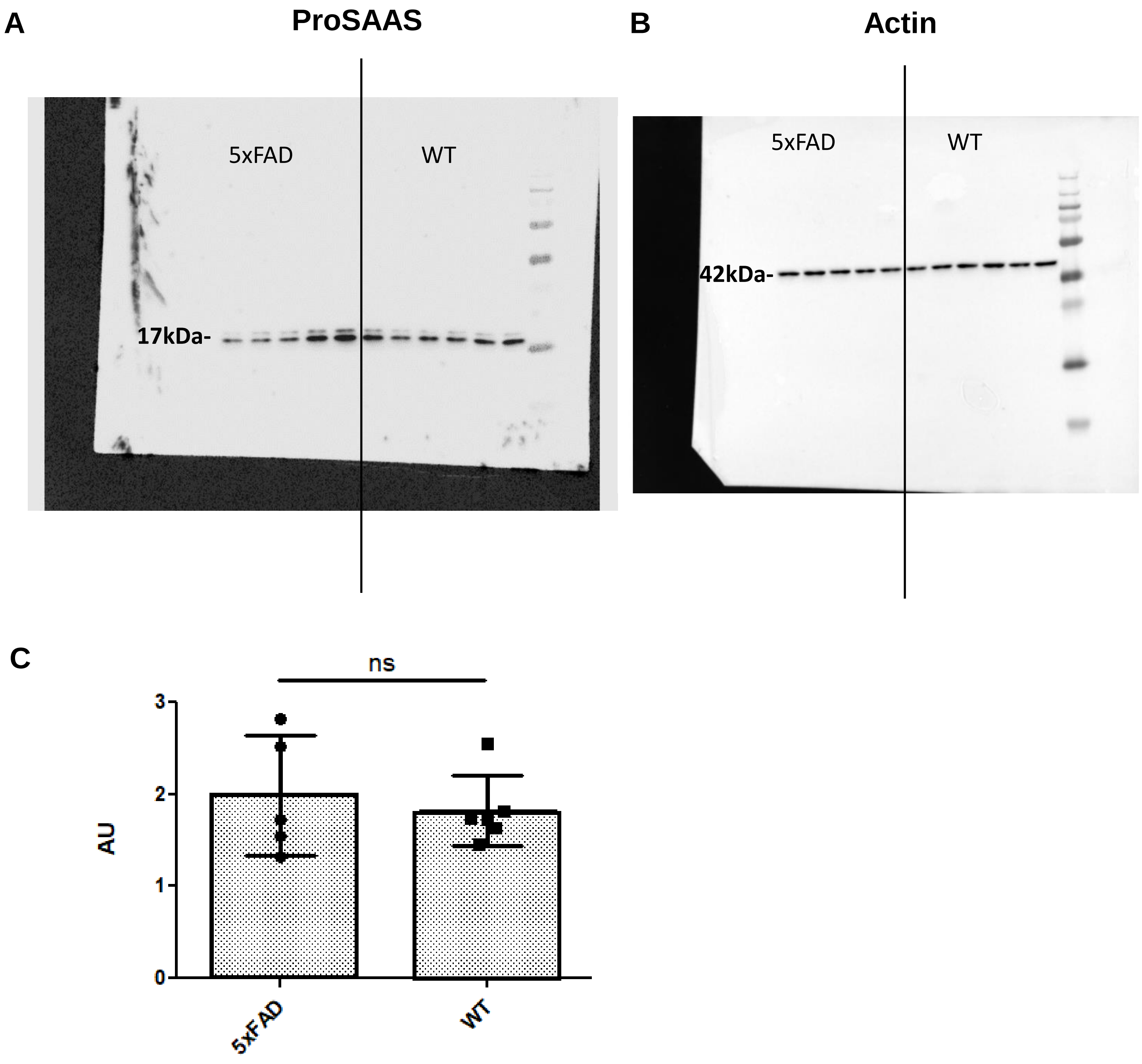


**Supp. Figure 6. Western blot of hippocampal extracts for proSAAS and actin shows no difference in proSAAS content between control and 5x FAD mice.** Hippocampi from WT and 5xFAD mice were extracted with detergent solution containing protease inhibitors, sonicated, centrifuged, and the clear supernatant was Western-blotted for proSAAS (**A**) and actin (**B**). Each band represents one mouse (both hippocampi were pooled). (**C**) Quantitation shows no significant differences between the proSAAS levels in the WT and 5xFAD mice extracts. Each data point represents the arbitrary value of band intensity from the proSAAS Western blot, normalized to the actin band in that sample. An unpaired t-test shows no significant differences.
